# Analysis of different strains of the turquoise killifish *Nothobranchius furzeri* identifies transcriptomic signatures associated with heritable lifespan differences

**DOI:** 10.1101/2023.11.20.567823

**Authors:** Mariateresa Mazzetto, Kathrin Reichwald, Erika Kelmer Sacramento, Philipp Koch, Alessandro Ori, Marco Groth, Alessandro Cellerino

**Affiliations:** BIO@SNS, Scuola Normale Superiore, Piazza dei Cavalieri,7, Pisa, 56126, Italy; Yale School of Medicine, 333 Cedar Street New Haven, CT 06510, USA; Leibniz Institue on Aging, Fritz Lipmann Institute, Beutenbergstr. 11, Jena, D-07745, Germany; Genetech DNA Way, South San Francisco, CA 94080, USA

## Abstract

The killifish *Nothobranchius furzeri* is the shortest-lived vertebrate model organism and is characterized by laboratory strains differing in lifespan by more than a factor of two. The genetic architecture underlying this difference is complex and the pathways responsive remain elusive. We performed an analysis in *N. furzeri* of public transcritomic datasets comprehensive of four different tissues and one embryonic stage in two strains: the shorter-lived GRZ and the longer-lived MZM0410. Remarkably, the two strains differ in their transcriptome profile already at the embryonic stage. The short-lived strain GRZ shows an anticipated aging profile consistently in all tissues investigated, with differences in expression of aging-related genes detected already at sexual maturity. In addition, analysis of a longitudinal dataset revealed that genes whose expression is prognostic of longer lifespan at the individual level are also differentially expressed between strains at an early adult age, suggesting antagonistic pleiotropism.

## 1| Introduction

Age-dependent transcript regulation has been investigated extensively in several models and in humans, yet the impact of genetic variability on age-dependent gene regulation is yet to be completely understood [1]. Differences in lifespan between individuals of the same species can be influenced by intrinsic- (random variations) and extrinsic-factors (such as nutrition) [2,3,4]. Disentangling the effects of genetic heterogeneity and stochasticity on lifespan of mammals is technically challenging and the majority of studies have been performed using invertebrates, such as the roundworm, *C. elegans* [4,5,3].

The annual African turquoise killifish (*Nothobranchius furzeri*) was recently established as a short-lived vertebrate model for aging research [6]. *N. furzeri* is particularly suited to investigate the effects of interventions on lifespan and aging-associated dysfunctions. Its maximum lifespan is 3-12 months, depending on the strain [7], the shortest lifespan ever recorded in captivity for a vertebrate [8]. Along with its short lifespan, the Turquoise killifish shows an accelerated aging phenotype, which shares many molecular and behavioral aging hallmarks that have been described in mammals, like accumulation of lipofuscin, gliosis, shortening of telomeres and cognitive decline [9].

A specific feature of this model is the large influence of the genetic background on lifespan: *N. furzeri* is indeed characterized by the presence of multiple laboratory strains whose founders originate from different spot habitats in Mozambique and Zimbabwe with large differences in captive lifespan [7] that can be as short as 3 months in the GRZ inbred strain [8] while recently wild-derived strains have lifespans in the order of at least 6-8 months and do not replicate GRZ extreme phenotype [10,11,12]. QTL mapping studies revealed a complex genetic architecture underlying this lifespan difference [13,14] that is therefore not the result of the fixation of a single deleterious mutation and must arise from a combination of naturally-occurring alleles fixed in the GRZ strain.

RNA sequencing (RNAseq) studies have demonstrated that the patterns of genome-wide transcript regulation in the *N. furzeri* mimic those observed in mammals [15,16] and this species was used in longitudinal experiment to investigate the correlation between individual lifespan and individual patterns of gene expression early in adult life [17,18].

In this study, we exploited *N. furzeri* transcriptomic dataset in order to identify gene expression signatures of genetically-determined lifespan differences and to compare these with patterns of age-dependent gene expression. We analyzed the impact of age and genotype on gene expression across different tissues. We then compared gene expression profiles of aging biomarkers in two strains and performed network analysis using weighted gene co-expression network analysis (WGCNA) [19], in order to correlate gene modules with external variables (strain in this case) and identified central hubs in the co-expression network of genes associated with heritable lifespan differences.

## 2| Methods

### 2.1| Transcriptome quantification and analysis

In general, sequencing of RNA samples was performed using Illumina’s next-generation sequencing methodology. In detail, total RNA was quantified and quality checked using Agilent 2100 Bioanalyzer Instrument (RNA 6000 Nano assay). Libraries were prepared from 500 ng of total RNA using TruSeq RNA v2 library preparation kit (Illumina) according the manufacturer’s instructions and subsequently quantified and quality checked using Agilent 2100 Bioanalyzer Instrument (DNA 7500 assay). Libraries were sequenced using a HiSeq 2500 System running in 51 cycle/single-end/high output mode. Sequence information was converted to FASTQ format using bcl2fastq v1.8.4. The RNA sequencing reads were aligned to the *N. furzeri* genome assembly (assembly version Nfu 20150522, gene annotation version 150922, both downloaded from the Nothobranchius furzeri Information Network Genome Browser) using STAR 2.5.1b (parameters: –alignIntronMax 100000, – outSJfilterReads Unique). For each gene, all reads that map uniquely to one genomic position were counted with FeatureCounts 1.5.0 under default settings [69]. Sequencing, mapping and counting qualities were assessed with MultiQC 0.9.

### 2.2| Proteome quantification and analysis

#### Sample Preparation

Individual brains from the fish were collected and snap frozen in liquid nitrogen. On preparation for MS, the brains were transferred into 1.5 mL Eppendorf tubes and 500 µL lysis buffer (4% SDS, 100 mM HEPES, pH8, 1mM EDTA, 100 mM DTT) added on ice. Samples were then vortexed (5 times) prior to sonication (Bioruptor, Diagenode), high setting, 20 °C, 10 cycles 1 min, with 30 sec interval). The samples were then centrifuged (3000 xg, 5min, room temp and the supernatant transferred to 2mL Eppendorf tubes. Reduction (15 min, 45 °C) was followed by alkylation (IAA 20 mM, 30min, RT, dark). Protein amounts were estimated, following an SDS-PAGE gel of 4% of each sample against an in-house cell lysate of known quantity. Between 200 and 300 µg of each sample was taken along for digestion. Proteins were precipitated overnight at 20 °C after addition of a 4x volume of ice cold acetone. The following day, the samples were centrifuged (30 min, 4 °C, 20800 xg) and the supernatant carefully removed. Pellets were washed twice with 1 mL ice-cold 80% acetone/20% water (10 min centrifugation, 4 °C, 20800 xg). They were then allowed to air-dry before addition of digestion buffer (3M Urea, 100 mM HEPES, pH8). Samples were resuspended with sonication (as above). LysC (Wako) was added at 1:100 enzyme:protein w:w and digestion proceeded for 4h at 37 °C with shaking (1000 rpm for 1h, then 650 rpm). Samples were then diluted 1:1 with MilliQ water and trypsin (Promega) added at the same enzyme to protein ratio. Samples were further digested overnight at 37 °C with shaking (650 rpm). The following day, digests were acidified by the addition of TFA (5%, 5 µ) and then desalted with Waters Oasis® HLB µElution Plate 30 µm (Waters Corporation, Milford, MA, USA) in the presence of a slow vacuum. In this process, the columns were conditioned with 3×100 µL solvent B (80% acetonitrile; 0.05% formic acid) and equilibrated with 3x 100 µL solvent A (0.05% formic acid in Milli-Q water). The samples were loaded, washed 3 times with 100 µL solvent A, and then eluted into PCR tubes with 50 µL solvent B. The eluates were dried down with the speed vacuum centrifuge and dissolved at a concentration of 1 µg/µL in 5% acetonitrile, 95% Milli-Q water, with 0.1% formic acid.

#### TMT labelling

The resuspended peptides (at 1 µg/µL) were rebuffered to pH 8.5 using 200 MM HEPES buffer (1:1 ratio) for labelling. 25 µg peptides were taken for each labelling reaction.

TMT-10plex reagents (Thermo Scientific) were reconstituted in 41 µL 100% anhydrous DMSO. TMT labeling was performed by addition of 2.5 μL of the TMT reagent. After 30 minutes of incubation at room temperature, with shaking at 600 rpm in a thermomixer (Eppendorf) a second portion of TMT reagent (2.5 μL) was added and incubated for another 30 minutes. After checking labelling efficiency by removal of 1 µg of material for MS analysis, samples were pooled (48 µg total), cleaned once again with Oasis and subjected to high pH fractionation prior to MS analysis.

#### High pH fractionation

The high pH fractionation was performed as described in Buczak et al (PMID: 32737464, [1]). Offline high pH reverse phase fractionation was performed using an Agilent 1200 Infinity HPLC System equipped with a quaternary pump, degasser, variable wavelength UV detector (set to 254 nm), peltier-cooled autosampler, and fraction collector (both set at 10 °C for all samples). The column was a Gemini C18 column (3 m, 110 Å, 100 1.0 mm, Phenomenex) with a Gemini C18, 4, 2.0 mm SecurityGuard (Phenomenex) cartridge as a guard column. The solvent system consisted of 20 mM ammonium formate (pH 10.0) as mobile phase (A) and 100% acetonitrile as mobile phase (B). The separation was accomplished at a mobile phase flow rate of 0.1 ml/min using the following linear gradient 1% B for 2 min, from 10% B to 40% B in 100 min, to 85% B in a further 1 min, and held at 85% B for an additional 5 min. We collected 48 fractions and pooled them to 16 samples for MS analysis. Pooled fractions were dried and resuspended in 0.1% formic acid and 5% acetonitrile for MS analysis.

#### Data Acquisition

Data acquisition has been performed using the settings described in Buczak et al. (PMID: 32737464, [1]) for SPS-MS3 acquisitions. In short, all acquisitions were done on an Orbitrap Fusion Lumos mass spectrometers (Thermo Fisher Scientific, San Jose, CA) using the Proxeon nanospray source and coupled to a nanoAcquity UPLC (Waters, Milford, MA) fitted with a trapping (nanoAcquity Symmetry C18, 5 μm, 180 μm × 20 mm) and an analytical column (nanoAcquity BEH C18, 1.7 μm, 75 μm x 250 mm). Solvent A was water, 0.1% formic acid and solvent B was acetonitrile, 0.1% formic acid.

Full scan MS spectra with mass range 375-1500 m/z were acquired in profile mode in the Orbitrap with resolution of 60,000 FWHM (at 200 m/z) using the quad isolation. The most intense ions from the full scan MS were selected for MS2, using quadrupole isolation. HCD was performed in the ion trap with normalized collision energy of 35%, with an intensity threshold of 5,000. For the MS3, the precursor selection window was set to the range 400–2000 m/z, with an exclude width of 18 m/z (high) and 5 m/z. The most intense fragments from the MS2 experiment were co-isolated (using Synchronus Precursor Selection = 8) and fragmented by HCD (collision energy, 65%). MS3 spectra were acquired in the Orbitrap over the mass range 100–1,000 m/z and resolution set to 30,000.

#### Data analysis

TMT data were processed using Proteome Discoverer v2.0 (Thermo Fisher Scientific). Data were searched against nothobranchius_furzeri (in-house) and a contamination database using Mascot v2.5.1 (Matrix Science) with the following settings: Enzyme was set to trypsin, with up to one missed cleavage. MS1 mass tolerance was set to 10 ppm and MS2 to 0.5 Da. Carbamidomethyl cysteine was set as a fixed modification and oxidation of methionine as variable. Other modifications included the TMT-10plex modification from the quan method used. The quan method was set for reporter ions quantification with HCD and MS3 (mass tolerance, 20 ppm). The false discovery rate for peptide-spectrum matches (PSMs) was set to 0.01 using Percolator. Reporter ion intensity values for the filtered PSMs were exported and processed using in-house written R scripts to remove common contaminants and decoy hits. Additionally, only PSMs having reporter ion intensities above 1 × 103 in all the relevant TMT channels were retained for quantitative analysis.

### 2.3| Differential expression analysis

Differentially expressed transcripts or miRNAs for the different comparisons were obtained with the DESeq2() package [20]: the differential expression results were obtained with alpha=0.05. DEGs were filtered for padj<0.05 and then used for visualization.

Differential expression analysis was applied on proteins using limma() package using normalized IBAQ values as inputs: differentially expressed proteins were filtered for padj<0.05 and used for visualization. Only protein groups quantified by at least two unique peptides were analyzed for differential expression. Data were analyzed using the MSnbase package. Reporter ion intensities were log2-transformed and normalized using the vsn package. Peptide-level data were summarized into their respective protein groups by taking the median value. Differential protein expression was assessed using the limma package. Differences in protein abundances were statistically determined using the Student’s t test moderated by the empirical Bayes method. P values were adjusted for multiple testing using the Benjamini Hochberg method.

### 2.4| Enrichment analysis

Gene Ontology was performed for each tissues with the WebGestalt online tool using GO Process; enriched categories for each quadrant were filtered for FDR<0.05. To combine the results metanalysis was performed using the Fisher’s Method, and then p-values were adjusted. Generally Applicable Gene-set/Pathway enrichment (GAGE) was performed using the gage() package [21]; enriched categories were filtered for q-value<0.05 and used for visualization with the Revigo software. Entrez IDs were used as identifiers for the analysis, from the human orthologues gene symbols: these were obtained with the biomart() package [22,23].

### 2.5| Principal Component Analysis (PCA)

Visualization of the data (for both transcriptome and miRNome) was performed with Principal Component Analysis (PCA): differentially expressed genes for different feature selection were obtained and selected for the analysis. Principal Components of the samples were obtained through the prcomp() function and then plotted, after computing centroids as means of the replicates for each group of samples.

### 2.6| Analysis of aging biomarkers

Transcripts with monotonic trend were used from [18]: only gene lists for cluster 1 and cluster 2 were considered for the analysis. Scaled expression values were computed after centering the raw counts to the mean and scaling on the standard deviation, and mean values for each time point were obtained. Visualisation of the results was performed using ggplot2() package [24].

### 2.7| Network analysis

Weighted gene correlation network analysis (WGCNA) analysis was performed using the WGCNA package in R [25] using the unsigned correlation as option and setting the minimum cluster size to 30 members. Input for WGCNA was a list of 4247 genes (with number of counts>100 for each sample): consensus analysis was performed, using the public pipeline, on brain, liver, skin and muscle. The selected module was then visualized using Cytoscape and filtering the network on the weight of connection (values>0.08).

### 2.8| Data and code availability

The accession numbers for the data reported in this paper are GEO: GSE52462 (MZM aging for brain, liver and skin), GSE103816 (MZM aging, muscle), PRJEB5837 (diapause and non-diapause, GRZ and MZM, embryo), GSE103132 (GRZ aging, brain), GSE103137 (GRZ aging, liver), GSE103140 (GRZ aging, skin), GSE103815 (GRZ aging, muscle), GSE92854 (small-RNAseq, both MZM and GRZ for brain, liver and skin), GSE207748 (validation dataset) GSE98744 (transcriptome for *H.glaber*, liver).

Codes are available upon request from the corresponding author.

### 2.9| Animal procedures and Ethical statement

Fish were bred at the Leibniz Institute on Aging, Fritz Lipmann Institute, Jena. Procedures for fish breeding, husbandry and euthanasia were performed in accordance with the rules of the German Animal Welfare Law and approved by the Landesamt für Verbraucherschutz Thüringen, Germany.

Eggs were maintained on wet peat moss at room temperature in sealed Petri dishes. When embryos had developed, eggs were hatched by flushing the peat with tap water at 16–18°C. Embryos were scooped with a cut plastic pipette and transferred to system tank. Fry were fed with newly hatched Artemia *nauplii* for the first 2 weeks and then weaned with finely chopped Chironomus *larvae*.

The system water temperature was set at a constant 27°C. At the desired age, fish were sacrificed via anesthetic overdose (Tricaine, MS-222) in accordance with the prescription of the European (Directive 2010/63/UE) and the rules of the German Animal Welfare Law.

## 3| Results

### 3.1| Genotype has a large effect on gene expression in *Nothobranchius furzeri*

In order to identify the transcriptional signature associated to heritable lifespan differences, we analysed 211 RNA-seq libraries obtained from two strains differing in lifespan (MZM0410 and GRZ), four tissues (brain, liver, skin, muscle) and 5 time points. The time points collected correspond to sexual maturity (5 weeks) and (for MZM0410) to young adult (12 weeks post hatching, wph), adult (20 wph), old (27 wph), and geriatric (39 wph), with the exception of muscle for which only young (9 wph) and old (27 wph) samples were available. For GRZ, the time points collected were 5 wph, 7 wph, 10 wph, 12 wph, and 14 wph. In addition, we analysed embryos at mid somitogenesis stage in two conditions (diapause and direct development) (Fig. 1A). To corroborate these results, we also analysed 150 libraries of miRNA-seq [26] obtained from a subset of the same adult samples.

**Figure 1.**
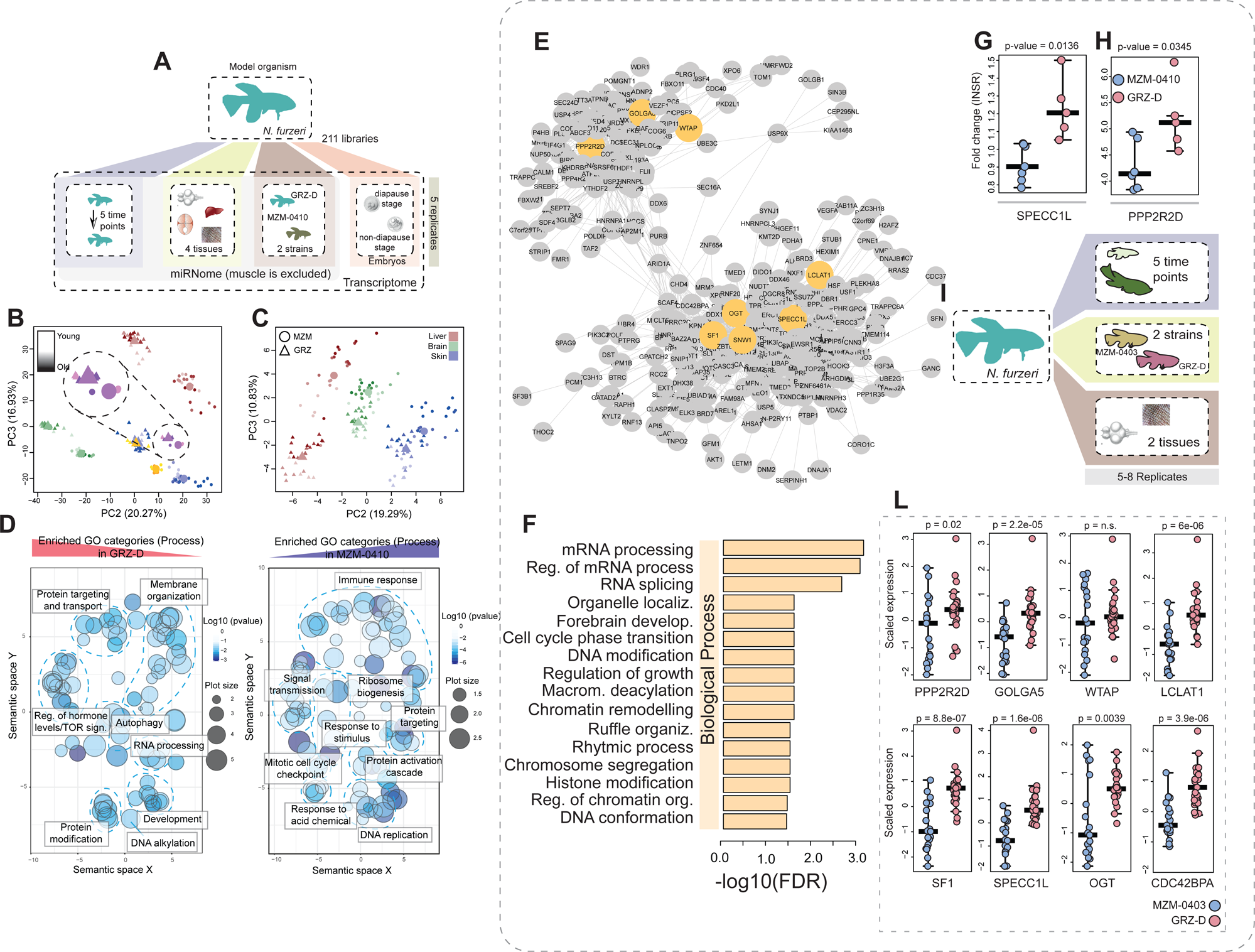
(A) Description of the discovery datasets: an RNA-seq dataset comprehensive of 5 tissues (brain, liver, skin, muscle and embryo), 2 strains (MZM-0410 and GRZ-D), 5 different time points (for brain, liver and skin) and 2 different stages (for embryo), with 5 replicates per sample and a small RNA-seq dataset (Baumgart, 2017) containing 3 different tissues (brain, liver and skin) and up to 6 different time points. (B) Principal Component Analysis (PCA) of brain (green), liver (red), skin (blue), muscle (yellow), and embryos (violet) samples based on transcript expression. Circles represent MZM samples, triangles represent GRZ samples; intensity of hues code for the time points. Inset shows a magnification of the embryo samples. (C) Principal Component Analysis (PCA) of brain (green), liver (red), skin (blue), muscle (yellow), and embryos (violet) samples based on miRNAs expression. Circles represent MZM samples, triangles represent GRZ samples; intensity of hues code for the time points. (D) Gene set enrichment analysis for strain-dependent transcripts. Enrichment analysis was performed with the gage() package and enriched categories were summarized and displayed using REVIGO (Supek, 2011). Significance of the single categories is coded by the intensity of the hue; GO terms were clustered according to semantic similarity and manually grouped into functional clusters (light blue dashed lines). (E) Overview of the “turquoise” gene module obtained by WGCNA: only the centre of the module, containing nodes with connection weight > 0.08, is visualised. Nodes highlighted in orange are the genes whose differential expression was tested in the validation set. Embryo samples were excluded from the network analysis. (F) Over-representation analysis of GO terms in the “turquoise” module obtained from WebGestalt: all Biological Process categories with FDR < 0.05 are shown. (G-H) Validation by qPCR of the hub genes SPECC1L (G) and PPP2R2D (H): qPCR was performed on MZM0410 and GRZ brain samples (5-weeks old) and Cq values were normalised to expression of INSR. Statistica, significance was determined with the Student t-test. (I) Description of the validation dataset. (L) Expression of the genes highlighted in (E) in the validation RNA-seq dataset. Statistical significance was computed with the Wilcoxon test. Only brain samples were considered for the analysis.

Principal component analysis (PCA) was used to visualize the transcriptional similarities between samples and revealed a clear separation based on the tissue, but also of samples from different strains within the same tissue. Interestingly the direction separating the two strains ws consistent across tissues (Figure 1B). Surprisingly, this separation was detected already at the embryo stage (Figure 1B, inset). The same tissue-independent effect of genotype on global expression patterns could be observed also for miRNAs (Figure 1C). Differential expression analysis was performed with DESeq-2 [20] using generalized linear models to separate the effects of different independent variables on gene expression. To identify gene sets that are differentially expressed in the two strains, we performed generally applicable gene set enrichment (GAGE) analysis [21] excluding the embryo samples. GAGE detected significant enrichment for a large number of gene sets. To facilitate visualization, these sets were further grouped into semantically similar clusters using REVIGO [27] (Figure 1D). Categories related to development, RNA processing, protein-targeting, -transport or -modification, autophagy and membrane organization were upregulated in the short-lived strain (GRZ-D). Genes upregulated in the long-lived strain were found to be enriched in categories related to DNA replication/cell cycle and immune response/protein activation cascade. These results highlight profound and unexpected differences in gene expression between these two strains.

Some genes showed extreme differences in expression between the two strains, for example the genes dead-end (DND), that is normally expressed during early embryonic development of the primordial germs cells [28] and in adult gonads, and the transporter SLC22A13, whose expression is normally restricted to the kidney [29], have negligible expression in MZM0410, but are strongly upregulated in all tissues of the GRZ strain (Figure S1). Examples of the opposite pattern are ACO2, coding for aconitase 2, and MACROD2, a gene whose deletion causes chromosome instability and represents a cancer risk [30] (Figure S1). In order to validate these findings, we exploited an independent RNA-seq dataset that contains samples related to two different strains at different time points: GRZ and MZM0403, a longer-lived strain genetically distinct from MZM04010 [7] (Figure 1I). The selected genes indeed show a similar expression profile during aging; DND and SLC22A13 are overall more expressed in the GRZ strain, on the other hand ACO2 and MACROD2 are more expressed in the MZM0403 strain (Figure S3 A-D) in the validation set.

### 3.2| Network analysis identifies genes related to splicing and protein modification as hubs of gene modules related to genotypic differences

Network analysis identifies modules of co-regulated genes and their respective hubs. We applied the weighted gene co-expression network analysis (WGCNA) method, which groups genes into discrete modules based on the co-expression patterns of their corresponding transcripts or proteins quantified as topological overlap in a network space where edges are defined by a power function of correlation of expression levels across samples [25]. WGCNA also computes the eigengene for each module as a summary vector of the coherent regulation of genes within a module, which can be then related to an external trait (in this case, strain or age). We performed a variant of the WGCNA method, the consensus analysis among different datasets, in order to identify modules (and hub genes) conserved across tissues (brain, liver, skin and muscle). We analyzed 4247 transcripts that were detectable in all tissues (with counts per sample > 100); from the analysis, we obtained 10 modules, and we computed the association with the “strain” variable for each. We focused our attention on the “turquoise” module (Figure 1E), which had the highest statistical association with strain (computed from Fisher’s meta analysis among each tissue correlation value). The module was shown to be enriched in mRNA processing, transport-related and histone modification categories (Figure 1F). In order to validate these results, we selected genes with the highest intra-modular connectivity and gene-trait significance, which are highlighted as orange points in Fig 1E and are related to splicing, (such as SNW1, SF1 and WTAP), and to maintenance of organelle structure (such as GOLGA5). The differential expression of SPECC1L (involved in the organization of the actin cytoskeleton) and PPP2R2D (a phosphatase involved in the control of mitosis entry and exit) were validated by qPCR using a third independent biological sample of MZM0410 and GRZ brains (Figure 1G-H). Differential expression of hub genes was also validated by exploiting the independent RNA-seq dataset described above (Figure 1L).

### 3.3| GRZ strain shows an anticipated aging expression profile

To investigate the relationships between the effects of age and strain on global gene expression, we analysed the transcripts that were differentially expressed (FDR< 0.1) both during aging in MZM0410 and between MZM0410 and GRZ at the first time point analysed (9 weeks for muscle, 5 weeks for the other three tissues) and plotted the log2(fold change) during aging on the X axis and the log2(fold change) GRZ vs. MZM0410 on the Y axis for each tissue separately. In all four tissues, the DEGs showed preferentially the same direction of regulation in the two conditions and the data points were therefore concentrated in quadrants I and III (i.e. genes down-regulated with age are also lower-expressed in GRZ and *vice versa*, Fisher’s exact test: pvalue < 2.2e-16 for all the 4 tissues) (Figure 2A-2D). Similar positive correlations between the effects of age and strain was obtained also when only differential expression according to age in MZM0410 was used as filtering criterion (Figure S2A-S2D).

**Figure 2.**
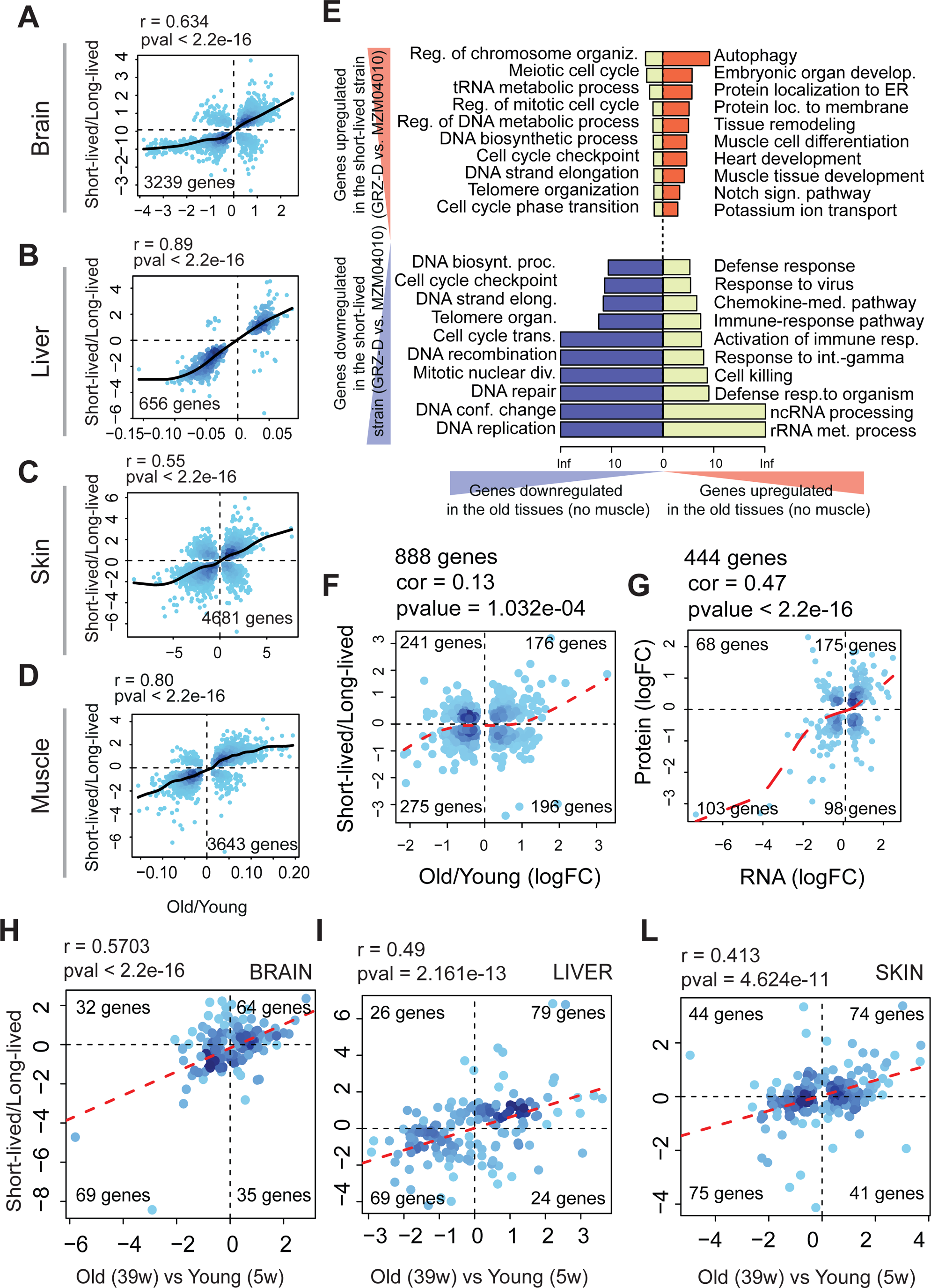
(A-D) Scatterplots of strain- and age-dependent DEGs in the transcriptome from A) brain, B) liver, C) skin and D) muscle. For each DEG, Log_2_ Fold change for Old vs. Young comparison is plotted on the X axis and Log2 fold-change Short-lived vs. Long-lived strain comparison on the Y axis. Only genes DEGs significant in both comparisons are shown. Pearson’s correlation coefficient and total number of DEGs in the intersection as reported on the graphs. The density of genes is coded by the intensity of the blue hues, the black line represents a spline fit of the data. (E) GO summary analysis of the genes contained in the 4 quadrants of the plots shown in Fig. 3A-3D. Gene enrichment was performed for each tissue separately using Biological Process as a the GO term database and then a metanalysis was performed using Fisher’s method. The length of the bars represents –log10 of the false discovery rate (FDR). (F) Scatterplots of age- and strain-dependent protein expression in the brain of *N. furzeri*. Log2 fold changes for age-regulated proteins are shown on the X axis and for strain-dependent proteins on the Y axis. Only proteins that are significant for both the comparisons are plotted. Pearson’s correlation and total number of genes, as well as number of genes per quadrant are shown. The density of genes is coded by the intensity of the blue hues. (G) Comparison between strain-dependent differentially-expressed transcripts and proteins in the brain of *N. furzeri*. LogFC values for transcripts are shown on the X axis and for proteins on the Y axis. Only genes significantly affected both at the transcript and protein level are plotted. Pearson’s correlation, total number of genes in the intersection, as well as number of genes per quadrant are reported. The density of genes is coded by the intensity of the blue hues. (H-L) Comparison between strain and age-dependent differential expression in the miRNome of brain (H), liver (I) and skin (L). Gene regulation is plotted as Log_2_FC for Old vs. Young comparison on the X axis and Short-lived vs. Long-lived strain comparison on the Y axis. MicroRNAs differentially expressed in either of the two conditions (union) are shown. Pearson’s correlation coefficient, total number of DEGs and number of genes per quadrant are reported. The density of genes is coded by the intensity of the blue hues.

In order to identify the biological processes that are differentially expressed according to strain and age, we performed GO over-representation analysis for the genes in the four quadrants of Figure 2A-D. Results of the three individual tissues were combined using Fisher’s metanalysis where the p-value of each GO represents a summary of the p-value for over-representation in the individual tests. Results are displayed in Figure 2E. Genes consistently down-regulated in the short-lived strain and during aging were enriched in terms related to cell cycle, DNA replication, cell-proliferation and -differentiation, indicating that age-dependent decrease in mitotic activity is anticipated in GRZ. Genes up-regulated in GRZ and during aging are enriched in terms related to autophagy, protein localization and I-kB/NF-kB signalling. Interestingly, genes up-regulated during aging and down-regulated in GRZ show a highly significant enrichment for the terms ncRNA processing and defence response. Genes down-regulated during aging and up-regulated in GRZ are enriched in terms related to cell cycle.

In order to validate these findings, we exploited the independent RNA-seq dataset described above and repeated the analysis shown in Figure 2A-D. In this case, we analysed the transcripts that were detected as differentially expressed both during aging in MZM0403 (comparing 287 days-old animals to 21 days-old animals) and between MZM0403 and GRZ at the first time point analysed (21 dph) and plotted the log2(fold change) for aging on the X axis and for strain on the Y axis (Figure S3E). Notably, the first time point of this analysis is two weeks earlier than in the previous analysis and precedes sexual maturity. In both comparisons, the DEGs showed preferentially the same direction of regulation in the two conditions (as observed for the GRZ vs. MZM0410 comparison) and the data points were therefore concentrated in quadrants I and III, with a strong positive correlation r = 0.604 for GRZ vs. MZM0403). The genes down-regulated in the GRZ strain and in old animals (for both comparisons) were enriched in categories related to genome maintenance and development (reproducing the results shown in Figure 2), while genes up-regulated for both the variables were enriched in rRNA metabolic process and protein localization to ER, but a significant enrichment of the autophagy term was not detected in this comparison (Figure S3F).

The correlation analysis was performed also on differentially expressed microRNAs, and it consistently showed a positive correlation between age- and stain-dependent regulation of miRNAs (Figure 2H-L).

### 3.4. Proteome analysis confirms differential expression results

In order to validate transcriptome regulation at the protein level, we performed mass-spectrometry-based proteomics on an independent sample of five young (5 weeks-old) GRZ brains and we compared strain-dependent expression at the transcriptomic and proteomic level. The results indicate a clear positive correlation between differentially expressed proteins and transcripts in the comparison of the two strains at 5 weeks (Figure 2F).

We then plotted age and strain-related differentially expressed proteins, as performed previously for the transcriptome (Figure 2A-D): we used adult (12 weeks-old) and young (5 weeks-old) brain samples from the long-lived strain for the aging comparison, and young GRZ vs. MZM0410 brain samples for the strain comparison. We detected a positive correlation between age- and strain-dependent protein regulation (Figure 2G).

The analysis done on transcripts and proteins was performed also on differentially expressed miRNAs, and it consistently showed a positive correlation between age- and stain-dependent small non coding transcripts (Figure 2H-L).

### 3.5. Cox-Hazard model reveals genes with antagonistic effects on lifespan that show a strain/age-dependent expression profile

To assess whether age- and strain-dependent transcript regulation is relevant for lifespan determination, we re-analyzed a longitudinal RNAseq dataset, where transcripts from fin biopsies were quantified at 10 and 20 wph, and individual lifespan information is available for each subject [17] (Figure 3A). We then applied the Cox-proportional-hazard model to this dataset. Cox analysis quantifies the effects of an independent variable, transcript levels in our case, on mortality as an exponential coefficient β. Negative values of log(β) indicate that the higher the gene expression, the lower the mortality risk and positive values of log(β) indicate that the lower the gene expression, the lower the mortality risk (Figure 3B). In the scatterplot in fig. 3B, the log(β) of individual genes at 10 wph and 20 wph are reported separately. The majority of significant cases are concentrated in quadrant II, with a log(β) of opposite signs at the two tested ages. Since factors with positive sign log(β) are risk factors and factors with negative sign log(β) are protective factors, this result implies that the expression at the majority of observed significant genes has an antagonistic action: beneficial at early time points and detrimental at later time points (quadrant II), or the opposite (quadrant IV).

**Figure 3.**
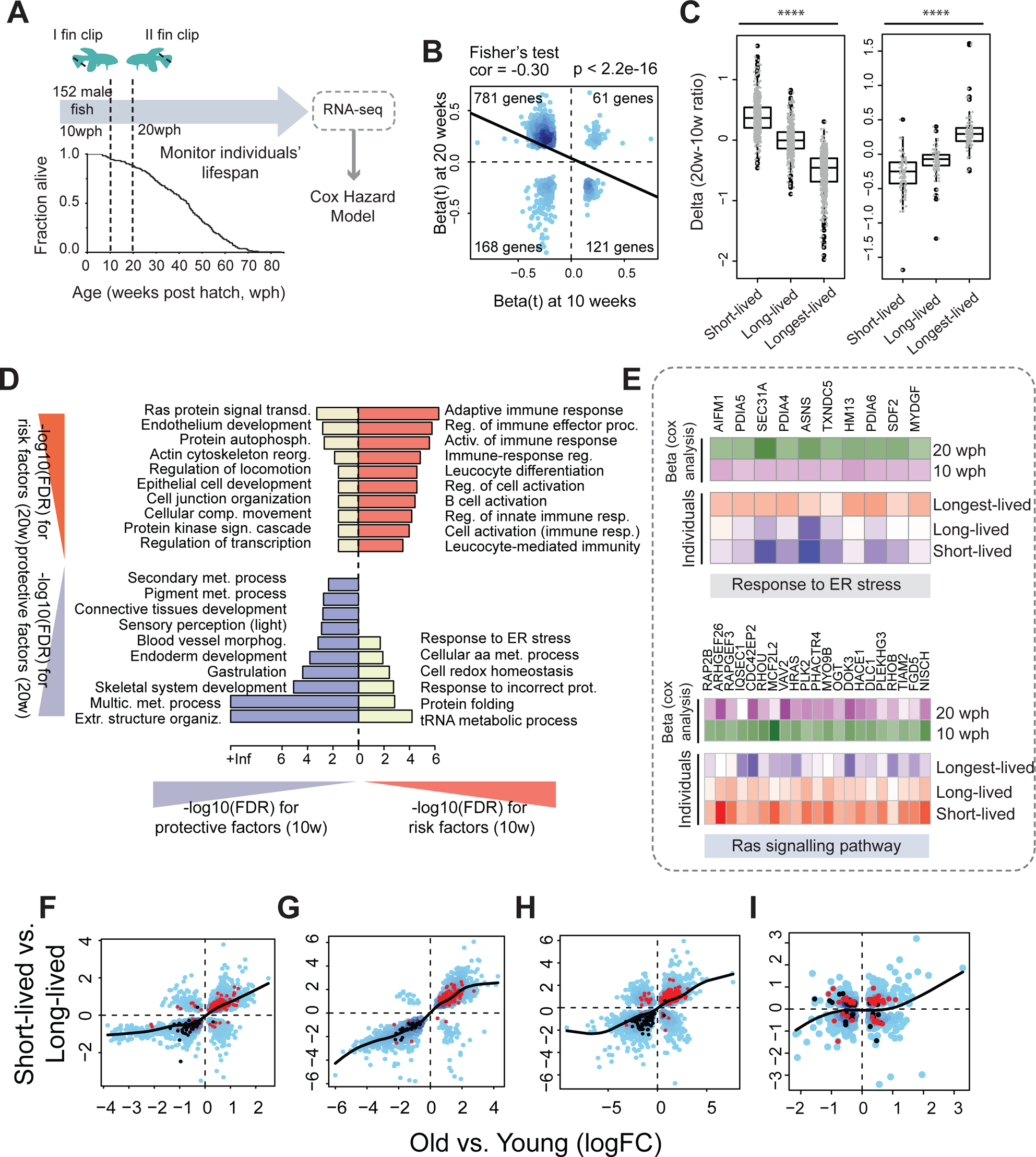
Cox regression analysis on a longitudinal dataset. (A) Workflow of the longitudinal experiment (modified from [15]). (B) Scatterplot beta coefficients obtained from Cox hazard regression analysis at 10 weeks (x-axis) and 20 weeks (y-axis). Only genes with significant beta coefficients for both the analyses (pval < 0.05) are displayed. Fisher’s test results and total number of genes, as well as number of genes per quadrant are shown. The density of genes is coded by the intensity of the blue hues. (C) Age-dependent regulation (shown as 20w/10w ratio) of genes with antagonistic pleiotropic action (i.e. genes contained in Quadrant II in the right-side plot and factors contained in Quadrant IV in the left-side plot) in different classes of individuals (from [15]): short-lived, long-lived and longest-lived animals were taken into account, as individuals with lifespans respectively between 28-36 wph (short-lived), 45-50 wph (long-lived) and 57-71 wph (longest-lived). Statistical analysis was performed with ANOVA test. (D) GO analysis of the genes contained in the 4 quadrants of the plot showed in Fig. 3B. Gene enrichment was performed using GO Process as a functional database; lenght of the bars represents –log10(FDR). (E) Detail on ER stress and Ras signalling pathway as examples of specific enriched categories with antagonistic pleiotropic action. Age-dependent regulation (as ratio 20w/10w) in short-lived, long-lived and longest-lived animals is displayed as gradient from red (upregulated) to white (neutral) to blue (downregulated), while beta coefficients (from Cox analysis) are displayed as gradient from purple (beta>0) to green (beta<0). (F-I) Aging- and strain-dependence of factors with antagonistic pleiotropic behaviour. Scatterplot of strain and age-dependent DEGs in (F) brain, (G) liver and (H) skin transcriptome, as well as in brain proteome (I) displayed as in Fig. 2. Genes in quadrant II and IV of Fig. 3B are highlighted as red and black dots, respectively.

We also analyzed the expression profile (as 20w-10w ratio) of genes in Quadrant II or IV for individuals in different classes of survivorship (Figure 3C): short-lived with age of death between 28 and 36 weeks, long-lived with age of death between 45 and 50 weeks and finally longest-lived with age of death between 57 and 71 weeks [15]. Genes contained in Quadrant II of Figure 3B increased expression on average with age in short-lived animals and decreased expression on average in the longest-lived animals; on the other hand, genes contained in Quadrant IV displayed decreased expression with age in short-lived animals and increased expression in longest-lived animals.

GO overrepresentation analysis was then performed on the genes contained in each quadrant of the plot (Figure 3D). Risk genes (quadrant I) were enriched in terms related to immune response (as expected, since upregulation of the immune system at old age can induce chronic inflammation with increased mortality) [31]. On the other hand protective genes (quadrant III) were found to be highly enriched in extracellular matrix organization (e.g. Collagen). Antagonistic genes with initial protective effect (quadrant IV) were enriched in response to ER stress, protein folding and homeostasis, and antagonistic genes with initial detrimental effect (quadrant II) were enriched for Ras protein signal transduction (quadrant II), as shown also when regulation of individual genes of these categories are displayed (Figure 3E): categories that have beneficial effect at early time points and detrimental effects at later time points show a progressive decrease in expression in long- and longest-lived animals (Ras signalling and protein phosphorylation) while categories that show the opposite effect on lifespan are progressive increased in long- and longest-lived individuals (response to ER stress).

To display the relationship between risk/protective factors from longitudinal datasets and cross-sectional datasets, we combined these two analysis: the genes shown in Figure 2A-C were displayed for brain, liver and skin and antagonistic genes were highlighted in different colors in these datasets (Figure 3F-H). From the plots, it is evident that antagonistic factors with a late detrimental effect (red dots) are preferentially downregulated both in old animals and in short-lived animals; on the other hand, factors with the opposite risk profile (black dots) are preferentially upregulated both in old and short-lived animals. These results could also be confirmed using the validation dataset presented in Figure 1I (Figure 3I).

### 3.6. Aging biomarkers reveal an “anticipated” aging profile in GRZ strain

To compare expression of putative aging biomarkers between the two strains, we used fuzzy c-means clustering to isolate transcripts with either negative or positive monotonic expression profile [17] in at least 2 tissues from the longer-lived strain. We opted for using the gene list of each cluster and plot the scaled expression value for each strain, as shown in Figure 3. Transcript expression profiles show that MZM biomarkers in GRZ have an “anticipated aging” profile. In particular, the expression level in the first time point (5 wph) was lower in GRZ strain for down-regulated biomarkers (Figure 4A-4B), but higher for upregulated biomarkers (Figure 3C-3D), whereas slopes are reduced in both cases. This data suggests that GRZ shows higher biological age at the earliest time point analyzed rather than a steeper slope of age-related change.

**Figure 4.**
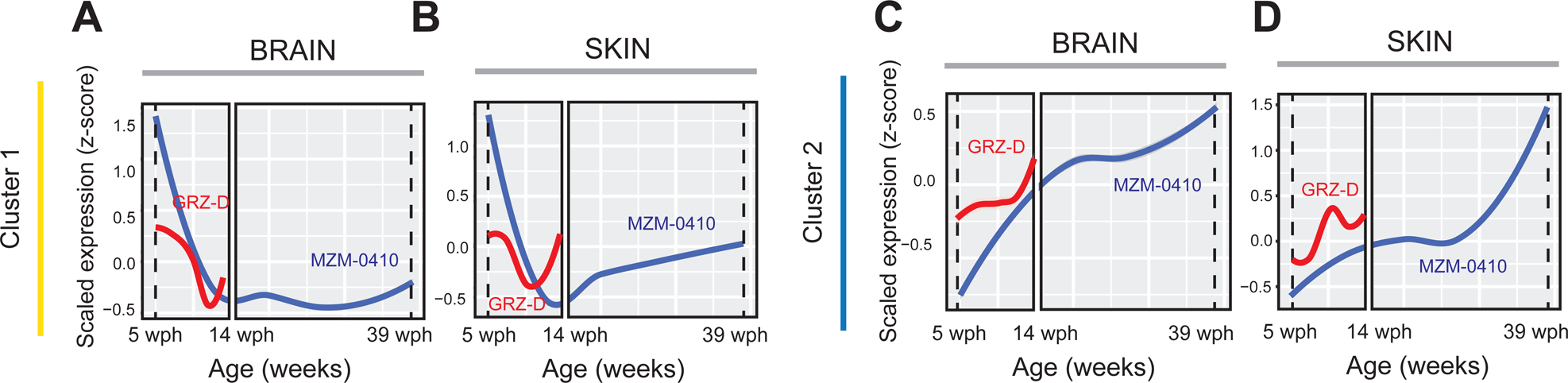
Comparison of expression of MZM-specific aging biomarkers in both strains (GRZ-D as red line and MZM0410 as blue line) for brain (A, C) and skin (B, D). The gene list for each cluster was obtained from [15]: clusters 1 (A-C) shows negative monotonic expression profile in all the analysed tissues, while clusters 2 (B-D) shows positive monotonic expression profile. Scaled expression values are plotted with the regression spline using the ggplot2 package, the shaded envelope represents confidence intervals of the mean expression.

### 3.7| Comparison with a long-lived model: the Naked Mole Rat

The previous results suggest that baseline transcript differences between short- and long-lived strains are correlated with age-dependent transcript regulation. A recent paper analyzed liver proteome of (*Heterocephalus glaber*), a notorious example of exceptional longevity [32] and guinea pig as a related shorter-lived species and detected a similar correlation between age-dependent expression and species-specific expression. Here, we analyzed a public dataset of age-dependent RNA-seq data from the skin and liver of naked mole rat [33] and compared these with RNA-seq data for the same organs of the guinea pig, which represents a shorter-lived close relative. Indeed, a recent paper analyzed liver proteome in these two species and revealed a correlation pattern between inter-species comparison and age-dependent expression similar to one we report in Figure 2A-2D. We intersected age-dependent DEGs in the naked mole rat with DEGs of the contrast between naked mole rat and guinea pig at young and displayed these as 2D plots of the log2(fold change) in either condition obtaining a significant positive correlation (Figure 5). The same result was obtained by analysing the union of the DEGs for skin (Figure 5A).

**Figure 5.**
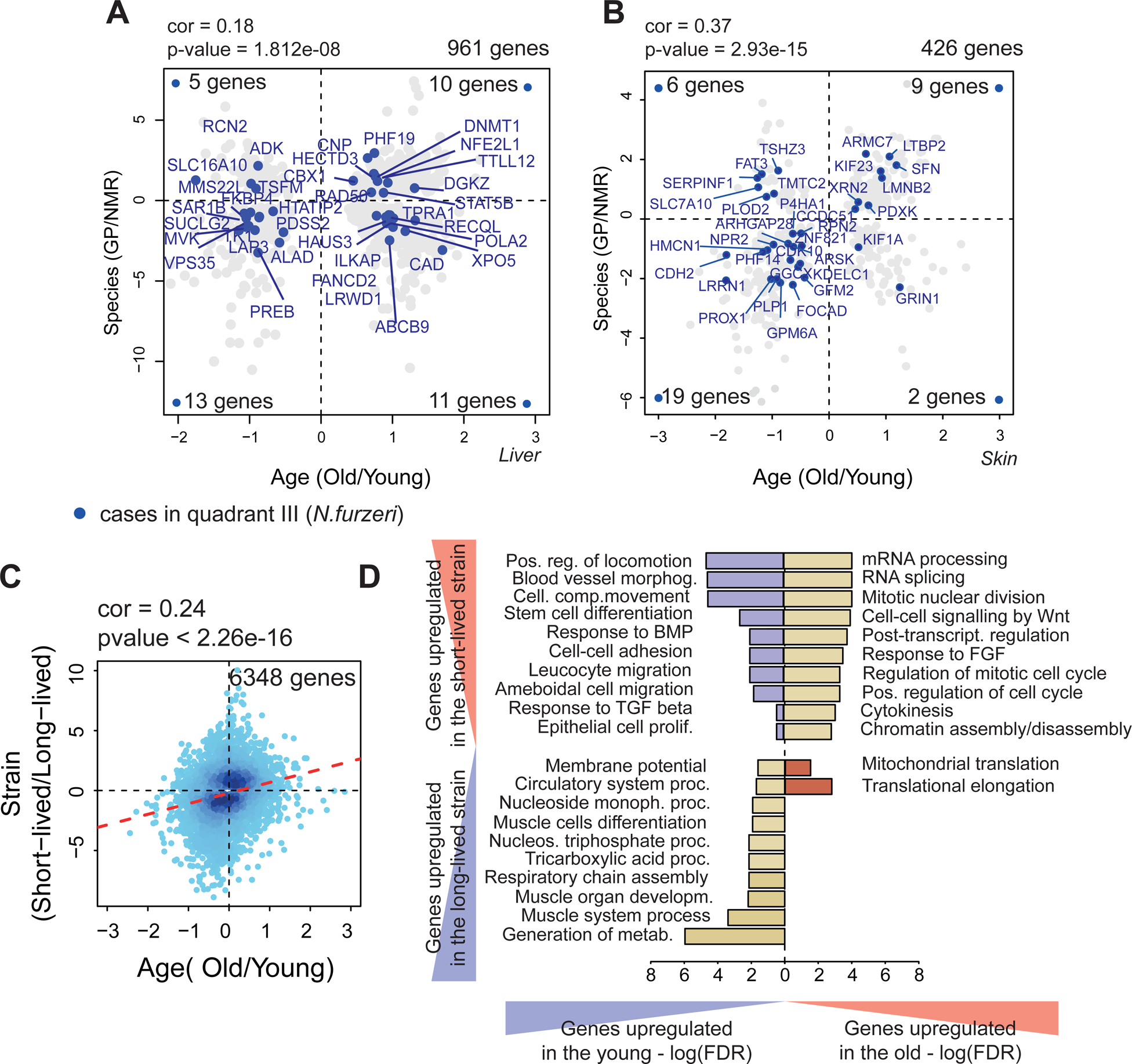
Comparison of age-dependent and lifespan-dependent gene expression between *N.furzeri* and *Heterocephalus glaber* in the liver (A) and in the skin (B). (A) Scatterplot of age-dependent DEGs in the Naked mole rat on the X axis and *H. glaber* vs. *Cavia porcellus* on the Y axis for liver samples (from [30]); LogFC for Old vs. Young and Long-lived vs. Short-lived comparisons are displayed. Only genes with padj < 0.01 for both comparisons are displayed. Pearson correlation and total number of significant genes are also shown in the plot. Coloured dots correspond to genes found in *N.furzeri* comparisons in the same tissue: in particular blue dots are genes downregulated in both the variables. (B) Scatterplot of age and strain DEGs in the Naked mole rat taken from skin samples; LogFC for Old vs. Young and Long-lived vs. Short-lived comparisons are displayed. Genes with padj < 0.01 for both the comparisons were considered as significant. Pearson correlation and total number of significant genes are also shown in the plot. Colored dots correspond to genes found in *N.furzeri* comparisons in the same tissue: blue dots are genes downregulated in both the variables. (C). LogFC value for Old vs. Young and Short-lived vs. Long-lived samples is displayed: filtering was performed on the combined pvalue between the two variables using meta-analysis. Pearson correlation and total number of genes, as well as number of genes per quadrant are shown. The density of genes is coded by the intensity of the blue hues. (D) GO analysis of the genes contained in the four quadrants of Fig. S7A. Gene enrichment was performed using GO Biological Process as a functional database. The categories are plotted as –log10 of the false discovery rate (FDR).

Further, we highlighted in Fig. 5A the genes that in *N.furzeri* were significantly downregulated both during aging and in GRZ strain (blue dots) (Figure 2A-2D). In the skin, these are concentrated in quadrant II, i.e. they are also downregulated during aging in the Naked mole rat and in the guinea pig (19 genes in quadrant III vs. 26 total, Fisher’s test pvalue = 2.45e-03, Figure 2, Figure 5C). The mentioned conserved genes (quadrant III in Figure 4B) are related to development and growth (such as PROX1 and NPR2), cell division/cytokinesis (such as HMCN1), neural stem cells differentiation and quiescence (such as GPM6A and CDH2), nervous system structure and development (such as PLP1 and LRRN1), and signal transduction (such as ARHGAP28).

In order to identify the biological processes that are differentially expressed according to species and age in the Naked Mole Rat, we performed GO over-representation analysis for the genes in the four quadrants of Figure S7A. The results (Figure 5D) show that genes down-regulated in the Guinea Pig and in old animals are enriched in categories related to membrane potential and metabolism. Interestingly, genes that are up-regulated during aging and in Guinea Pig show a highly significant enrichment for RNA processing and splicing (Figure S5A). These results are consistent with the prominent enrichment for the same terms in the gene coexpression network of the killifish transcriptome.

## 4| Discussion

We have analyzed the impact of genetically-determined lifespan differences on gene expression in the short-lived teleost model *Nothobranchius furzeri*, taking advantage of the large differences of lifespan among laboratory strains of this species and in particular the shorter-lived GRZ strain and the longer lived MZM0410 strain. To validate our results, we also analyzed a novel dataset where the GRZ strain is compared with the longer-lived MZM0403 strain.

One of the surprising results of our investigation is that the transctriptome patterns of the GRZ strain is clearly separated from that of the longer-living strains with genes that differ over 100-fold in expression. In addition, a difference in the transcriptome of the two strains can be detected already at the stage of somite embryos. The strain-specific signature is similar across organs as demonstrated by PCA. This baseline difference is overlaid onto differences in age-dependent regulation of gene expression that are consistent across tissues. This difference is detected as a positive correlation between difference in transcript abundance between strains at young and age-dependent differences in the longer-living strain. Genes whose expression is upregulated in the shorter-lived stain at young age also increase in expression with age and vice-versa. This result demonstrates that the profiles of the two strains diverge well before sexual maturity, suggesting an “anticipated” aging in the shorter-lived strain rather that an “accelerated” aging where the differences between strains only appears after sexual maturity and become progressively larger. One of the main results of this paper is indeed that the strain-specific transcript regulation can be observed during the early phases of development: genetic differences impact gene expression already at embryonic stage and seem to show their biggest effect at 5 weeks post hatching (corresponding to sexual maturity). On the other hand, all these changes appear smaller already at young adult stage (12 wph, following MZM-0410 timeline), and at that time point shorter-lived individuals show a slower aging rate, as compared to the longer-lived comparison strain. Analysis of monotonically regulated genes, which we consider as putative aging biomarkers, provides an independent support for this view. The anticipated aging process is associated to enrichment in autophagy (referred mostly to genes related to mitochondrial degradation) for up-regulated genes and cell cycle-related categories for down-regulated genes. Age-dependent decrease in cell cycle and development and upregulation of autophagy categories has been extensively reported in senescent cells [35,36].

An interesting correlation emerges also between genes whose expression is associated with lifespan differences within a strain (prognostic genes) and genes whose expression is associated to lifespan differences across strains. In particular, the largest proportion of prognostic genes show an antagonistic behaviour: high expression at young age is prognostic of longer lifespan but high expression at older ages is associated with higher mortality risk. These genes are also preferentially up-regulated in old animals and in the GRZ strain. These convergent lines of evidence suggest that upregulation of these genes is inducing damage and is reducing lifespan in the shortest-lived individuals.

The observation that higher expression of these genes has opposite association with mortality risk depending on age is also consistent with the *pleiotropic antagonism* theory positing that one gene controls multiple traits, where positive selection for an allele conferring advantage at young age is associated with detrimental phenotypes later in life. So an allele that increases reproduction in early life and induces faster late life senescence would be selected.

A specific pathway that emerged as particularly significant from our analysis is RNA splicing: network analysis revealed a conserved module across brain, liver, skin and muscle, which is positively correlated with genetic differences and is highly enriched in this term. Comparison of age and genetic-dependent DEGs in a long-lived species, the Naked Mole Rat (*Heterocephalous glabrus*), highlights again this category as up-regulated during aging in the naked mole rat and down-regulated in the Guinea Pig.

Other results reported in literature also point to a central role for splicing during aging: in the longitudinal work performed on *Nothobranchius furzeri* and presented by [17], the spliceosome complex was found among the top enriched categories in longest-lived animals. In monkeys, on the other hand, a direct link between nutrition and splicing was observed: caloric restriction (CR) showed to regulate RNA processing and to direct alternative splicing usage in the liver (Rhoads et al., 2018); in the transcriptome of human fibroblasts spliceosome was found to be downregulated with aging [35], consistently our previous cross-sectional work on *Nothobranchius furzeri* brain aging at the transcriptome - [15] and proteome-level [18].

Taken together, these results show that a genetic background which favours a shorter lifespan is often associated with an anticipated trasncriptomic aging signature, and that alterations in the splicing process can be affected by genetic differences, but that they can also be predictive for aging in longer-lived species.

## 6| Acknowledgements

The authors thank the fish facility of the Leibniz Institute on Ageing for support in the animal breeding and husbandry.

## 8| Tables

**Table S1:** GAGE output on strain-dependent DEGs.

**Table S2:** Pairwise comparisons age/strain in brain, liver, skin and muscle (with annotation).

**Table S3:** RNA/proteome comparison.

**Table S4:** Aging proteome.

**Table S5:** Pairwise comparisons age/strain in brain, liver, skin (with annotation) for miRNAs.

**Table S6:** Naked Mole Rat results.

**File S1.** R Markdown document with the computational pipeline.

**Supplementary Figure 1.** (A-D) Expression profile of some of the most DEGs between the two strains: DND (A), SLC22A13 (B), ACO2 (C) and MACROD2 (D). Normalized pseudo-counts were used for plotting expression strength; each point represents an individual case, the line connects the medians of each age group, MZM-0410 (orange) and GRZ (blue).

**Supplementary Figure 2.** A-D) Scatter plots of aging-dependent fold-changes in MZM-0410 (39 wph vs. 5 wph) vs. strain-dependent fold-changes in the young (5 wph). Only DEGs for the first comparison (Old vs. Young) are plotted and DEGs were computed for each tissue separately. The density of genes is coded by the intensity of the blue hues, the number of genes in each quadrant is reported in the respective vertex. (A) Brain, (B) Liver, (C) Skin and (D) Muscle.

**Supplementary Figure 3.** A-C) Scatter plots of aging-dependent fold-changes in MZM-0410 vs. aging-dependent fold-changes in GRZ. Only DEGs for the first comparison (Middle-aged vs. Young) are plotted and DEGs were computed for each tissue separately. The density of genes is coded by the intensity of the blue hues, the number of genes in each quadrant is reported in the respective vertex. (A) Brain, (B) Liver and (C) Skin. D-F) Scatter plots of strain-dependent fold-changes in the young (5 wph) vs. strain-dependent fold-changes in the middle-aged (12 wph). Only DEGs for the first comparison (Young GRZ vs. Young MZM) are plotted and DEGs were computed for each tissue separately. The density of genes is coded by the intensity of the blue hues, the number of genes in each quadrant is reported in the respective vertex. (D) Brain, (E) Liver and (F) Skin.

**Supplementary Figure 4.** (A) Heatmap of splicing-related genes in *Heterocepalous glabrus* and *Nothobranchius furzeri*. Only genes significant for both the comparisons and in both species are displayed. Upregulated genes (with logFC>0) are coloured as red boxes while downregulated genes (with logFC<0) are displayed as blue boxes.

