## Supplementary figures and images for "Analysis of different strains of the turquoise killifish *Nothobranchius furzeri* identifies transcriptomic signatures associated with heritable lifespan differences"

### Figure S1

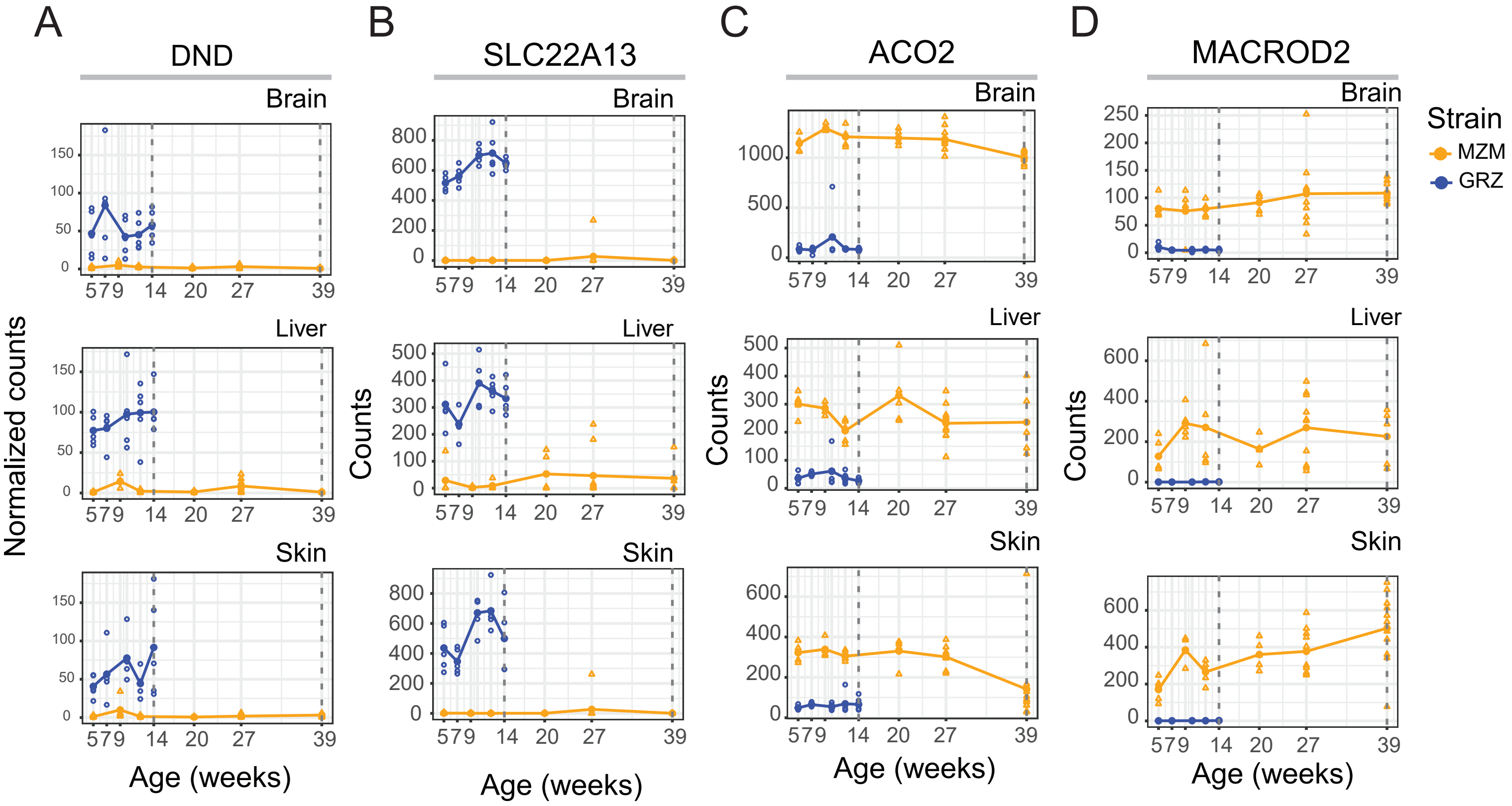

### Figure S2

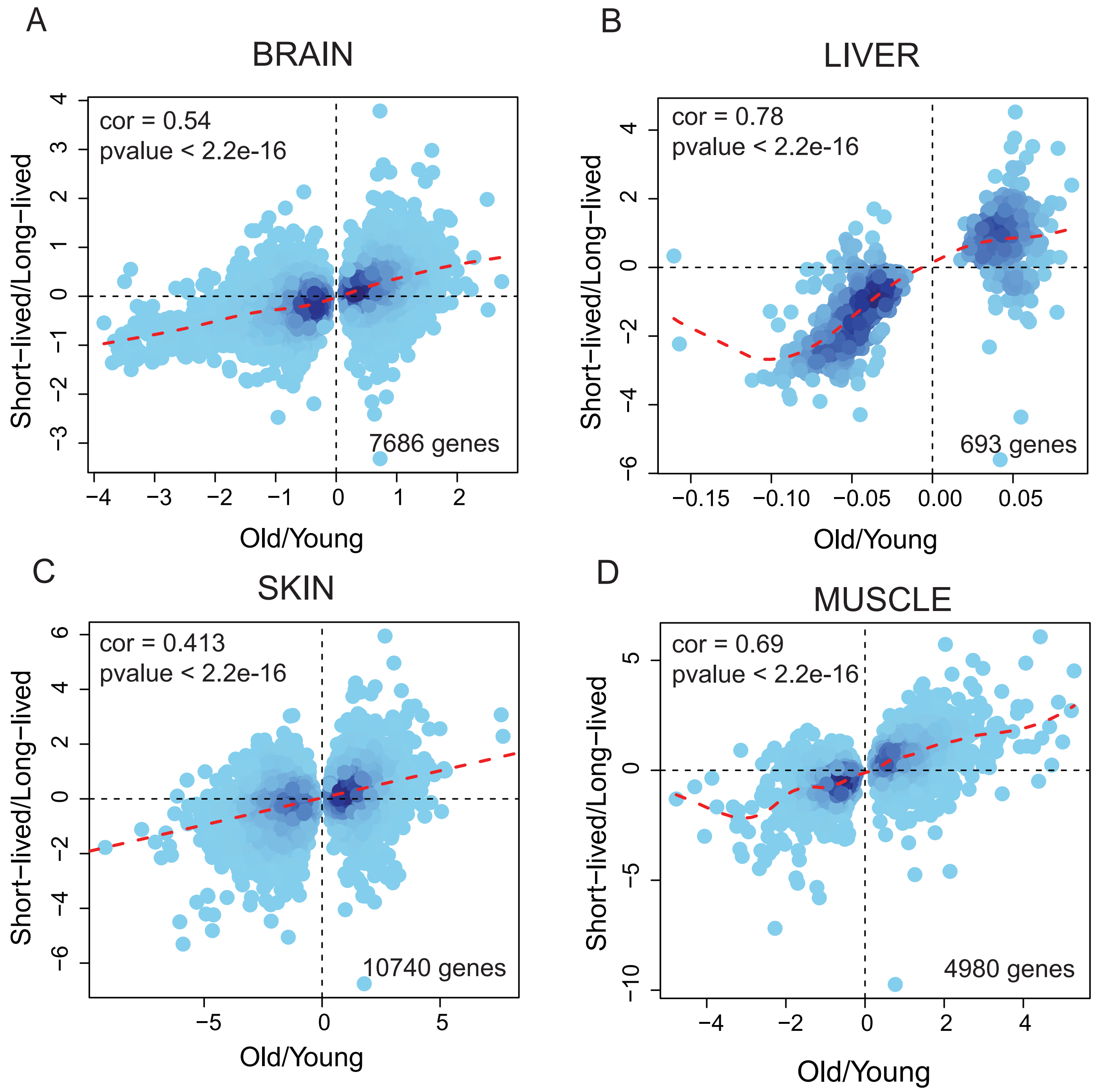

### Figure S3

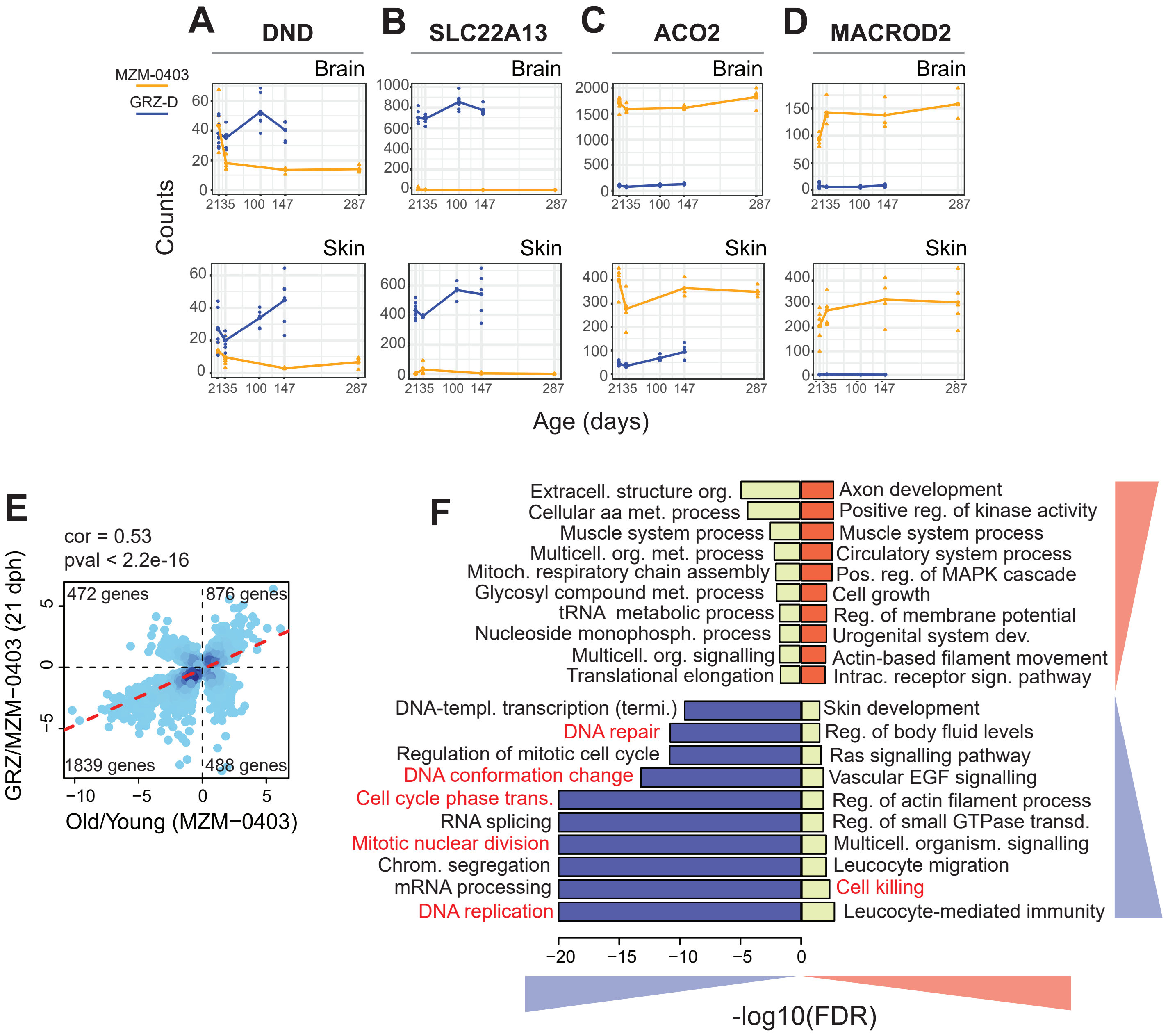

### Figure S4

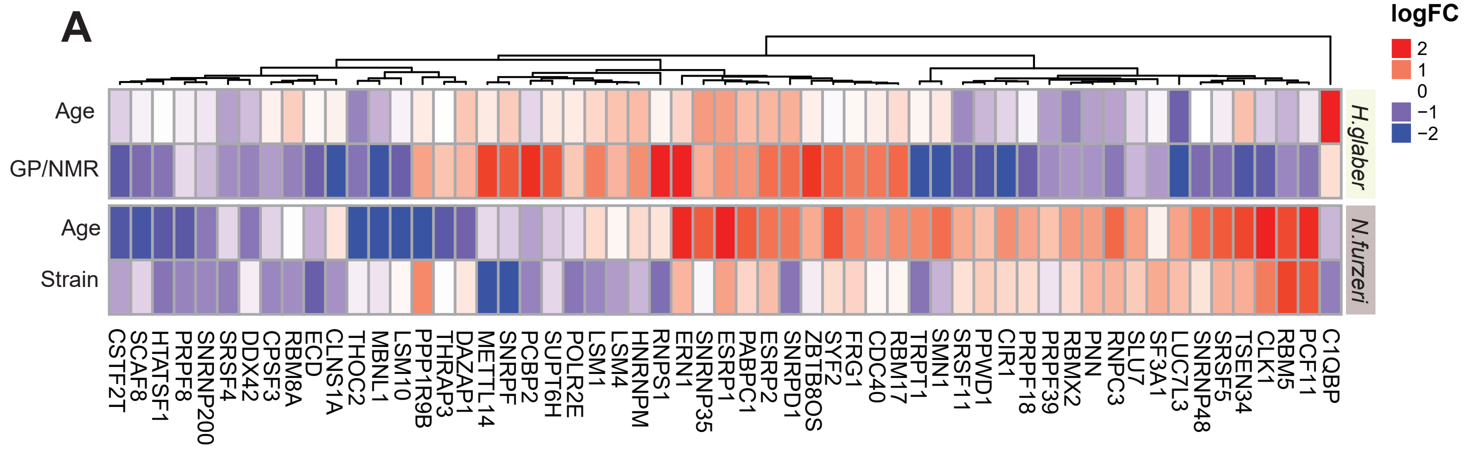
